# *sedimix*: A workflow for the analysis of hominin nuclear DNA sequences from sediments

**DOI:** 10.1101/2025.02.28.640818

**Authors:** Jierui Xu, Elena I. Zavala, Priya Moorjani

## Abstract

**Summary:** Sediment DNA––the ability to extract DNA from archaeological sediments–– is an exciting new frontier in ancient DNA research, offering the potential to study individuals at a given archaeological site without destructive sampling. In recent years, several studies have demonstrated the promise of this approach by recovering hominin DNA from prehistoric sediments, including those dating back to the Middle or Late Pleistocene. However, a lack of open-source workflows for analysis of hominin sediment DNA samples poses a challenge for data processing and reproducibility of findings across studies. Here we introduce a snakemake workflow, *sedimix*, for processing genomic sequences from archaeological sediment DNA samples to identify hominin sequences and generate relevant summary statistics to assess the reliability of the pipeline. By performing simulations and comparing to published studies, we show that *sedimix* has high sensitivity and precision. *sedimix* offers a reliable and adaptable framework to aid in the analysis of sediment DNA datasets and improve reproducibility across studies.

**Availability and Implementation:** *sedimix* is available as an open-source software with the associated code and user manual available at https://github.com/jierui-cell/sedimix

*Contact:* Jierui Xu

*Supplementary information:* Supplementary data are available at Bioinformatics online

## 1. Introduction

The ability to recover ancient hominin DNA from sediments[1–4] and other non-hominin skeletal remains[5] has revolutionized the study of human evolution. Sediment DNA has the potential to fill in gaps left by a sparse skeletal record and allow for more continuous explorations of human evolution through time and space. However, analyzing sediment DNA is inherently complex, as DNA from many diverse taxa (ancient and modern) are expected to be present, compounding the existing challenges of fragmented, deaminated sequences and low endogenous content inherent with ancient DNA[6–9]. The identification of authentic ancient hominin sequences is, thus, challenging and available pipelines for genetic analysis from metagenomic sources are not optimized for the identification and evaluation of ancient hominin DNA (e.g. *HOPS* and *MALT*). To our knowledge, there are no open-source workflows available for processing sediment sequencing data for the identification of hominin sequences, hindering the analysis and reproducibility of findings across studies.

Sediment DNA studies[3,4] utilize two primary approaches for the identification of hominin DNA sequences from metagenomic datasets. The first, composition-based methods, assign taxonomic labels to sequencing reads by comparing the data to existing databases of sequences from diverse species. These approaches use a *k*-mer based taxonomic classification using tools such as *Kraken2[10]* and *Centrifuge[11]*. Another approach, alignment-based methods, map the sequencing reads to multiple selected reference genomes. For alignment-based classification, both general-purpose aligners such as *bwa[12]* and *Bowtie2[13]* and specialized aligners suited for metagenomic datasets (i.e. *MALT[14]*) are used. For analysis of sediment DNA, it is typical to combine both alignment and classification steps, however, the order, the choice of methods and filtering steps differ and can impact the sensitivity and accuracy of identifying hominin DNA, as well as computational tractability of the analysis. Here we present a snakemake workflow, referred to as *sedimix*, for processing sediment DNA data to identify hominin DNA sequences. As input, *sedimix* takes raw sequencing data (*fastq* files) and returns an alignment file (*bam*) with sequences inferred as deriving from hominin and other sources. It also outputs a table of summary statistics per sample including the number of mapped sequences, average duplication rates, percent deamination, and percent hominin derived sequences. The user also has the option to generate a fastq file of all non-hominin sequences identified that may be useful for other analyses. We perform simulations to test the sensitivity, precision, and computational requirements of different steps of the workflow and apply *sedimix* to published datasets to demonstrate its reliability. *sedimix* offers a reliable and adaptable pipeline for the analysis of nuclear hominin sequences from archaeological sediment DNA datasets.

## 2. Materials and methods

Our workflow, *sedimix*, can be broken down into four main steps: filtering, classification, mapping, and generating summary statistics (**Figure 1**). In order to run the workflow the user must input (1) the *fastq* file(s) containing the raw sequencing reads, (2) the human reference genome sequence (in *fasta* format), and (3) a list of parameters to use (see *sedimix* github for details, https://github.com/jierui-cell/sedimix). The latter contains a list of parameters such as mapping quality and read length filters, identification of non-hominin reads, and selected classifier method (*Kraken2* or *Centrifuge*). In addition, the user can add an optional *bed* file containing a list of positions matching a SNP panel for capture array or ascertainment scheme of interest (in the same coordinates as the human reference genome used in input (2)). In this case, the reference genome will be modified at the specified positions to a non-reference or alternative allele in order to reduce reference bias as proposed in earlier studies[3,15].

**Figure 1:**
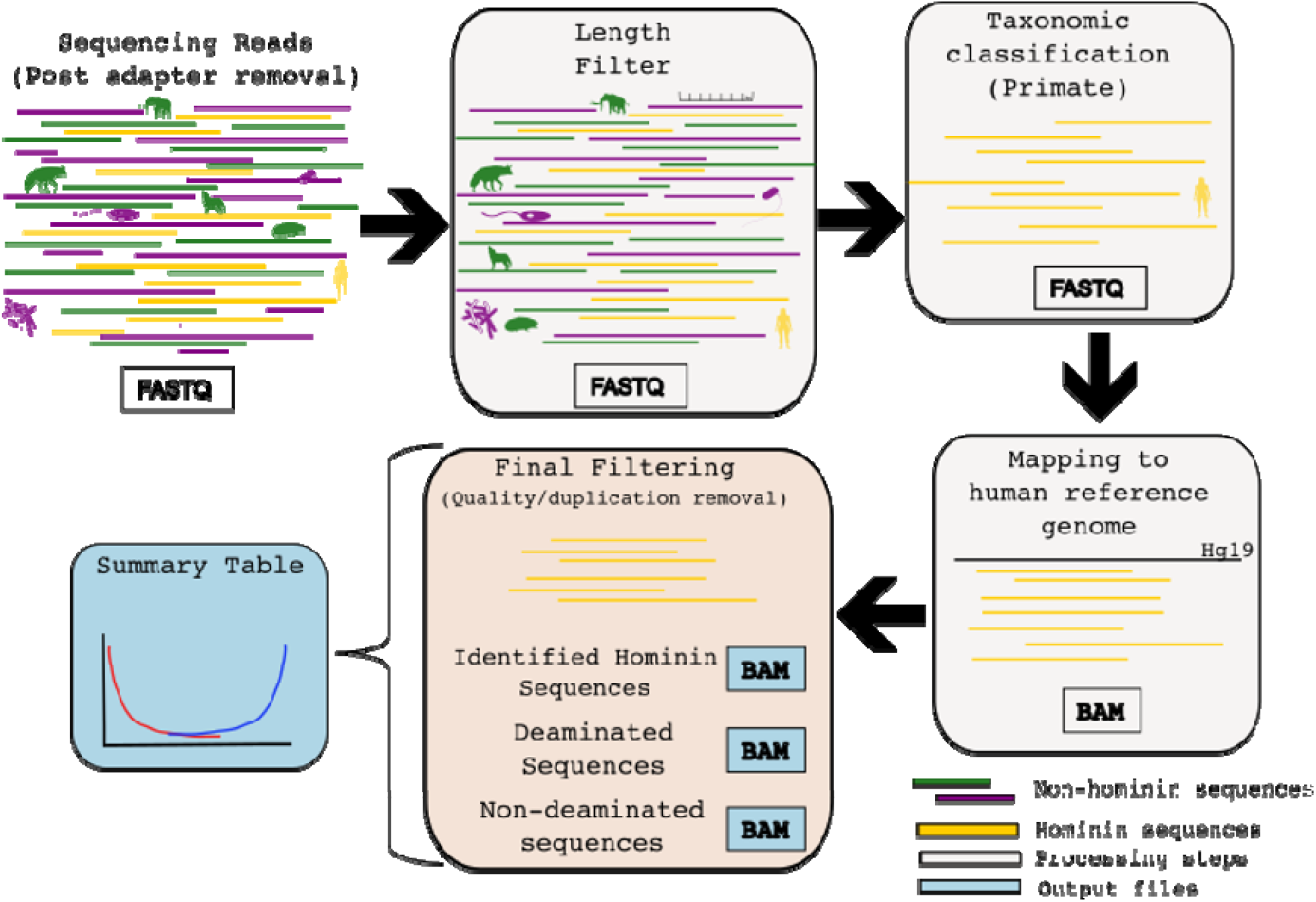
Overview of the different steps in the *sedimix* workflow. The input is a fastq file post adapter removal, which is then filtered based on read length. Next classification is performed and only reads identified as primate are retained. These reads are then mapped to the human reference genome followed by duplicate removal and filtering bas on mapping quality. The output are three different bam files and a table with summary statistics per fastq file processed.

*sedimix* workflow starts with the filtering step that involves removal of reads that are below a given length cut-off as specified in the parameter list (default: 30 base pairs). Next, the filtered reads are input into the specified classifier (default: *Centrifuge*, see **SI 3** for comparison of classifiers) and all the reads identified as ‘primate’ are extracted. While our goal is to identify hominin reads, we retain all sequences classified as ‘primate’ to maximize sensitivity at this step. Users can optionally extract reads classified to non-primate lineages for downstream analysis.

The third step is to map the primate reads to the user provided human reference genome using *bwa aln* with optimal parameters for ancient DNA specimens(options: -n 0.01 -o 2 -l 16500 [16]). Following mapping, sequences below a minimum mapping quality score (default: 25) and duplicates identified using *samtools markdup*[17,18] are removed. If a SNP panel was provided by the user, we use *bedtools intersect*[19] to subset the sequences that overlap the target SNP positions. This output *bam* file contains filtered sequences that map to the human reference genome. To support quality control analysis of these sequences, we split the output file into two separate files: one containing deaminated sequences which are a hallmark of authentic ancient DNA sequences (with a C-to-T or G-to-A substitutions in the first or last 3 bases) and the another one containing non-deaminated sequences. The last step in the workflow is to generate a report with summary statistics about the output bam file. The report contains the number of reads/sequences remaining after each filtering step, the number of deaminated sequences, the inferred percentage of deamination based on *mapdamage*[20], and an estimate of the percentage of hominin DNA within the primate sequences (see **SI2.4** and **Supplementary Table 2** for detailed descriptions).

## 3. Results

To evaluate the performance of *sedimix*, we simulated three datasets using an ancient DNA simulator, *gargammel[21]*. The simulated datasets had different compositions of sequencing reads as follows: (1) reads were composed of 90% modern human (Individual NA12778 from 1000 genomes[22]) and 10% Neanderthal (Altai[23]); (2) 40% modern human, 10% Neanderthal and 50% bacteria (composition from [24]); and (3) 10% modern human, 1% Neanderthal, 44.5% mammals and 44.5% bacteria(**Supplementary Table 3**). In total 10 million reads were generated for each simulated dataset. We added double-stranded deamination for the Neanderthal reads under the Briggs model[6] with an average length of 0.4 of single-stranded overhanging ends, 3% single-stranded nick frequency, 30% deaminated cytosine residuals in single-stranded DNA, and 1% deaminated in double-stranded DNA. We evaluated the performance by measuring two metrics: *sensitivity* defined as the number of identified reads that were correctly classified as ‘hominin’ (Neanderthal or modern human) divided by the total number of hominin reads simulated, and *precision* defined as the number of identified reads that were truly hominin divided by the total number of identified reads.

Across simulations, we find that *Centrifuge* consistently had higher sensitivity than *Kraken2*, with minimal impacts on precision and lower memory requirements (**SI 3.1**). Thus, we recommend *Centrifuge* as our default classifier. In addition, we find that performing classification before mapping reduces run times (**SI 3.2**) and using a classification cut-off at ‘primate’ rather than ‘*Homo sapiens’* increases the sensitivity of identifying hominin reads (**SI 3.3**). Finally, we show that precision is impacted if using a smaller reference database because there is higher mis-classification of reads from non-hominin mammals as primate/ hominin(**SI 3.4**). These tests inform our recommendations for the *sedimix* workflow, which focuses on enhancing computational tractability and retention of accurately classified hominin sequences.

Using these recommendations, we then tested the sensitivity and specificity (with default settings) of *sedimix*. For the three simulated datasets described above, we find *sedimix* has both high sensitivity and specificity. Specifically, we achieved a sensitivity of 94.1%, 94.1% and 94.7%, and a precision of 100%, 100%, 97.5% respectively for three simulated datasets. The run time increased linearly with the number of input reads, and processing 100 million reads took 45 minutes on 32 Intel Xeon Gold 6330 CPUs. The memory requirement measured as the maximum resident set size (RSS) was 139 GB throughout the workflow, which should be manageable on most virtual computers.

After validating *sedimix’s* performance on simulated datasets, we explored its performance in real data by using the sequencing data published in two recent studies. The first by *Gelabert et al*. included ancient human DNA that was found in a 25,000 year old sediment sample[4]. This dataset consists of a single *fastq* file with 522,997,582 raw sequencing reads. It required 4 hours and 5 minutes for the analysis and reached a maximum memory usage of 141 GB. We identified 0.1% hominin DNA with ∼20% deamination rates on both ends, in close concordance with previous findings (**SI 4**). The second dataset from *Essel et al*. included ancient human touch DNA that was recovered from a cervid tooth[5]. This dataset comprised five files with a total of 86,847,334 raw reads and took 2 hours and 3 minutes for the analysis, with a peak memory usage of 139 GB. We identified ∼1% hominin sequences across three files, which is slightly higher than the published estimates though our estimated deamination rates are similar. This analysis showcases the reliability of *sedimix* for the recovery of hominin nuclear sequences from archaeological sediment samples, providing a robust and flexible tool for the future metagenomics applications.

## Acknowledgements

J.X. was supported by the BWF grant to P.M. E.I.Z.’s work was funded by the Miller Institute in Basic Research Science at University of California, Berkeley and the Novo Nordisk Hallas-Møller Emerging Investigator Grant (NNF24OC0088862). P.M. was supported by NIH R35GM142978, Burroughs Wellcome Fund (Career Award at the Scientific Interface) and the Koret-UC Berkeley-Tel Aviv University Initiative in Computational Biology and Bioinformatics.

## References

1. Slon V, Hopfe C, Weiß CL et al. Neandertal and Denisovan DNA from Pleistocene sediments. Science 2017;356:605–8.

2. Zavala EI, Jacobs Z, Vernot B et al. Pleistocene sediment DNA reveals hominin and faunal turnovers at Denisova Cave. Nature 2021;595:399–403.

3. Vernot B, Zavala EI, Gómez-Olivencia A et al. Unearthing Neanderthal population history using nuclear and mitochondrial DNA from cave sediments. Science 2021;372, DOI: 10.1126/science.abf1667.

4. Gelabert P, Sawyer S, Bergström A et al. Genome-scale sequencing and analysis of human, wolf, and bison DNA from 25,000-year-old sediment. Curr Biol 2021;31:3564–74.e9.

5. Essel E, Zavala EI, Schulz-Kornas E et al. Ancient human DNA recovered from a Palaeolithic pendant. Nature 2023;618:328–32.

6. Briggs AW, Stenzel U, Johnson PLF et al. Patterns of damage in genomic DNA sequences from a Neandertal. Proc Natl Acad Sci U S A 2007;104:14616–21.

7. Allentoft ME, Collins M, Harker D et al. The half-life of DNA in bone: measuring decay kinetics in 158 dated fossils. Proc Biol Sci 2012;279:4724–33.

8. Sawyer S, Krause J, Guschanski K et al. Temporal patterns of nucleotide misincorporations and DNA fragmentation in ancient DNA. PLoS One 2012;7:e34131.

9. Dabney J, Knapp M, Glocke I et al. Complete mitochondrial genome sequence of a Middle Pleistocene cave bear reconstructed from ultrashort DNA fragments. Proc Natl Acad Sci U S A 2013;110:15758–63.

10. Wood DE, Lu J, Langmead B. Improved metagenomic analysis with Kraken 2. Genome Biol 2019;20:257.

11. Kim D, Song L, Breitwieser FP et al. Centrifuge: rapid and sensitive classification of metagenomic sequences. Genome Res 2016;26:1721–9.

12. Li H, Durbin R. Fast and accurate short read alignment with Burrows–Wheeler transform. Bioinformatics 2009;25:1754–60.

13. Langmead B, Salzberg SL. Fast Gapped-Read Alignment With Bowtie 2 Nature Methods. 2012; 9: 357--9.

14. Herbig A, Maixner F, Bos KI et al. MALT: Fast alignment and analysis of metagenomic DNA sequence data applied to the Tyrolean Iceman. bioRxiv 2016, DOI: 10.1101/050559.

15. Günther T, Nettelblad C. The presence and impact of reference bias on population genomic studies of prehistoric human populations. PLoS Genet 2019;15:e1008302.

16. Schubert M, Ginolhac A, Lindgreen S et al. Improving ancient DNA read mapping against modern reference genomes. BMC Genomics 2012;13:178.

17. Danecek P, Bonfield JK, Liddle J et al. Twelve years of SAMtools and BCFtools. Gigascience 2021;10, DOI: 10.1093/gigascience/giab008.

18. Bonfield JK, Marshall J, Danecek P et al. HTSlib: C library for reading/writing high-throughput sequencing data. Gigascience 2021;10, DOI: 10.1093/gigascience/giab007.

19. Quinlan AR, Hall IM. BEDTools: a flexible suite of utilities for comparing genomic features. Bioinformatics 2010;26:841–2.

20. Jónsson H, Ginolhac A, Schubert M et al. mapDamage2.0: fast approximate Bayesian estimates of ancient DNA damage parameters. Bioinformatics 2013;29:1682–4.

21. Renaud G, Hanghøj K, Willerslev E et al. gargammel: a sequence simulator for ancient DNA. Bioinformatics 2017;33:577–9.

22. 1000 Genomes Project Consortium, Auton A, Brooks LD et al. A global reference for human genetic variation. Nature 2015;526:68–74.

23. Prüfer K, Racimo F, Patterson N et al. The complete genome sequence of a Neanderthal from the Altai Mountains. Nature 2014;505:43–9.

24. Seguin-Orlando A, Korneliussen TS, Sikora M et al. Paleogenomics. Genomic structure in Europeans dating back at least 36,200 years. Science 2014;346:1113–8.

