## Supplementary Material for "*sedimix*: A workflow for the analysis of hominin nuclear DNA sequences from sediments"

#### 1. Introduction

*sedimix* presents a workflow for the identification of ancient hominin sequences from metagenomic sequencing data (i.e. environmental or sediment DNA). With this pipeline we aim to aid in the analyzing of sediment DNA samples and enhance reproducibility across studies. To assess the reliability of the pipeline, we performed simulations and chose optimized parameters to enhance the reliability while ensuring computationally tractability (**Supplemental Section 2 and 3**). We then applied the pipeline to two previously published datasets[1,2] (**Supplemental Section 4**). Below, we detail the different steps in the pipeline and the testing that was performed to examine the performance of the pipeline.

#### 2. Workflow Steps

*sedimix* was designed for identifying ancient hominin DNA (aDNA) from sediment sequencing data or other metagenomic datasets. Our pipeline consists of four steps: 1) filtering of reads, 2) classification (assigning reads specific taxa labels) with published methods such as Centrifuge, 3) mapping (aligning reads to the human reference genome) with *bwa* to identify human sequences, and 4) generating summary statistics (**Figure 1**). Below we describe each step in detail.

##### 2.1 Set-up and Input

*sedimix* is a snakemake workflow that should be run within a conda environment. Instructions on how to set-up the conda environment, including a file with the related python and R packages can be found on *sedimix*'s github (https://github.com/jierui-cell/sedimix). New users need to download several different software and classifier databases (**Supplementary Table 1**) before running *sedimix* for the first time. Details on how to complete these installations can be found on *sedimix* github. The locations of each of these software should also be added to the user's PATH. Users can test if the set-up was completed successfully by running "check_dependencies.py".

In order to run *sedimix*, users must provide (1) one or multiple fastq files containing genomic sequences from a sediment DNA sample, (2) the fasta file for the human reference genome, and (3) optionally a SNP panel containing a list of positions to restrict the analysis to. The fastq file should be demultiplexed (all reads should be from a single library) and the adapters should be trimmed.

If the user inputs a SNP panel bed file, it is recommended to generate an alternative human reference genome using the "generate_alternative_ref.py" script where the nucleotide sequence at the positions in the SNP panel are changed to a non-reference or alternative allele in order to minimize reference bias. It has been shown previously that otherwise reads carrying the reference allele will preferentially map, potentially leading to biases in downstream analyses [3,4]. The script assumes the bed file uses 1-based genomic position and is in the same coordinates as the reference genome being used.

**Supplementary Table 1:** Software and databases required for running *sedimix*. The version numbers cited are what was used by *sedimix* developers. Other versions may also work, but have not been tested. *Additional packages also required are included in the yaml file when building the *sedimix* conda environment.

| **Software/Data** | **Version** | **Reference/Relevant link(s)** |
|---|---|---|
| Centrifuge (workflow default) | 1.0.4 | [5] |
| Kraken2 (only if using this classifier) | 2.1.3 | [6] |
| Seqtk | 1.4 | [7] |
| BWA | 0.7.17 | [8] |
| Samtools | 1.14 | [9,10] |
| Bedtools | 2.28.0 | [11] |
| mapDamage | 2.2.1 | [12] |
| Human Reference Genome | hg19 | [13] http://hgdownload.cse.ucsc.edu/goldenPath/hg19/chromosomes/ |
| Python* | 3.11.6 | |
| R* | 4.4.0 | |
| Centrifuge NCBI non-redundant database | NCBI: nucleotide non-redundant sequences, Date: March, 2018 | https://benlangmead.github.io/aws-indexes/ |

| | | |
|---|---|---|
| Kraken2 NCBI non-redudant database (only required if Kraken2 is used instead of Centrifuge) | nt Database, Date: 11/29/2023 | https://benlangmead.github.io/aws-indexes/ |

#### 2.1 Filtering

Our workflow starts by performing filtering based on read length (default: 30 base pairs (bp)). Filtering based on read length is common in the processing of ancient and modern sequencing data as short reads have been shown to increase the likelihood of mismapping[14]. In ancient DNA analyses, this length is often restricted to 30 or 35 bp in an attempt to balance the retention of endogenous reads, while limiting errors from mismapped sequences[15]. We chose to start with this length filtering step, as performed in other workflows[1] to decrease the number of reads processed in the classification and mapping steps, therefore likely decreasing *sedimix's* run time.

#### 2.2 Classification

The next step involves classification of the length filtered sequences to different taxonomic groups in order to decrease faunal misassignment in nuclear sediment DNA. For this step, users can select either *Centrifuge* or *Kraken2*. *Kraken2* analyzes all k-mers of the same length in a read while *Centrifuge* starts at a segment with minimum exact match and extends as far as possible. We recommend *Centrifuge* due to its reduced memory requirement and comparable speed and accuracy to Kraken2 (**Supplemental Section 3.1**) . In addition, we also recommend using the NCBI nucleotide non-redundant sequence database to decrease potential faunal misclassification (**Supplementary Figures 5 & 6**).

#### 2.3 Mapping

The reads that are classified as "primate" are then extracted for subsequent genome mapping. We map the reads to the human reference genome using *bwa aln* with the optimal parameters for ancient DNA samples (-n 0.01 –o 2 –l 16500). After mapping, we remove duplicates and low mapping quality sequences (default: 25). If the user has provided a SNP panel, we only retain reads that overlap these positions. The resultant sequences are output as a bam file that contains sequences inferred as 'hominin' DNA. We then split this file into two additional files: deaminated hominin sequences and non-deaminated hominin sequences. The deaminated sequences are those that contain a C-to-T or G-to-A substitutions within the first three or last three base pairs. We acknowledge that expectations for the observations of deamination will be impacted by how libraries were prepared (i.e. single vs double-stranded library preparation). This definition is intended to be more general. However, if users prefer to use a different method or their own script(s), they can do so directly on the original identified hominin sequences BAM file after *sedimix* has finished running.

#### . 2.4 Summary Statistics

*sedimix* outputs a report file in tsv format containing multiple summary statistics per row for every input fastq file. The report file includes the number of mapped sequences, average duplication rates, percent deamination, and percent hominin derived. The latter is inferred by creating a list of lineage informative sites from previously published alignments of modern and archaic primates[16]. These sites were selected as described previously[3], where the Vindija33.19 Neanderthal, Altai Neanderthal, Denisova 3, human reference genome hg19, all individuals in 1000 genomes[17], pan troglodytes-2.1.4 (panTro4), and pan paniscus (panpan1.1) shared the same allele, which is different from that of pongo abelii (ponabe2) and macaca mulatta (rhemac3). Ponabe2 and rhemac3 must also share the same allele. We provide this file, but users are also able to input their own if they prefer. The different columns included in the report file are described in **Supplementary Table 2**.

**Supplementary Table 2**: Descriptions of the components in *sedimix*'s output report file.

| **Column Label** | **Description** |
|---|---|
| Sample ID | Extracted from input sequence file name. |
| Sequenced reads | Total number of input reads per sample (pre-filtering) |
| Length filtered | Total number of reads that pass the minimum length filter (default: 30 bp) |
| After classification | Total number of reads that are classified as Primate |
| Mapped sequences | Total number of sequences after mapping (pre-quality filtering and deduplication) |
| On-target sequences | Total number of sequences that overlap SNPs in provided bed file (this will equal mapped sequences if no bed file is provided) |
| Quality filtered | Total number of sequences that pass the minimum mapping quality filter (default: 25) |
| Unique filtered | Total number of sequences that are identified as unique (based on using samtools markdup -r) |
| Average duplication rate | Number of unique sequences divided by number of quality filtered sequences. |
| Number of SNPs covered | Total number of SNPs in the provided bed file that were covered by unique filtered sequences. This will be NA if no bed file is |

| | |
|---|---|
| | provided |
| Percent deaminated [%] | Percent of unique filtered sequences that are deaminated among all identified hominin sequences. |
| Percent hominin derived [%] | Percent of unique filtered sequences covering lineage informative sites that have the hominin allele |
| Percent hominin derived deaminated [%] | Percent of deaminated unique filtered sequences covering lineage informative sites that have the hominin allele |
| 5' C-to-T substitution frequency [%] (95% CI) | Percent of unique filtered sequences that have a T where the reference is a C at the 5' end, calculated from mapdamage2[12] output. |
| 3' C-to-T substitution frequency [%] (95% CI) | Percent of unique filtered sequences that have a T where the reference is a C at the 3' end, calculated from mapdamage2[12] output. |

### 3. Testing with Simulated Data

#### . 3.1 Comparison of Kraken2 and Centrifuge

Previous metagenomic workflows targeting the recovery of hominin nuclear DNA[1,3] have used either *Kraken[18]* or *Centrifuge[5]* for classifying reads by taxa. In order to select a classifier for our workflow, we performed two sets of simulations. First, we replicated a previous study comparing classifiers using viral genomes[19] as a sanity check. Second, we repeated the same comparison with varying complexities of metagenomic data containing ancient hominin DNA.

*Simulations with viral genomes*

While our focus is not on viral genomes, these comparisons are still useful and more computationally efficient in understanding larger trend differences between the different methods. Arizmendi Cárdenas et al.[19] used simulated viral genomes to compare the precision and sensitivity of *Centrifuge[5]*, *Kraken2[6]*, *DIAMOND[20]*, and *MetaPhlAn2[21]*. They found that *Centrifuge* and *Kraken2* had the best performance among these classifiers due to their ability to handle shorter reads (under default settings), recommending *Centrifuge* in their conclusions. We repeated their simulations using an ancient DNA simulator, *Gargammel[22]* to simulate sequencing reads from 238 reference viral sequences from 233 different human DNA viruses (one virus represented in six distinct contigs within RefSeq)[23–26]. These sequences were simulated for read lengths ranging from 30 bp to 150 bp, increasing in increments of 10 bp, for 10x coverage across each genome. We also simulate reads with increasing deamination from 0%

to 50% with 5% increment at read length of 60bp. For running *Kraken2*, we utilized the pre-built standard database that includes RefSeq archaea, bacteria, viral, plasmid, human, and UniVec sequences (78GB, https://genome-idx.s3.amazonaws.com/kraken/k2_standard_20240605.tar.gz). For *Centrifuge*, we also used the pre-built standard database available online that is similar to *Kraken2*, but is missing UniVec sequences (7.9GB, https://genome-idx.s3.amazonaws.com/centrifuge/p%2Bh%2Bv.tar.gz). For both classifiers, we used the default parameters recommended by the developers. We evaluated the performance for both methods by measuring the sensitivity (number of correctly identified reads divided by the total simulated reads) and precision (number of correctly identified reads divided by the total identified reads).

We find that *Centrifuge* outperforms *Kraken2*, with a higher average sensitivity and precision across the 238 simulated datasets (**Figure S1A & B**), aligning with the observations from Arizmendi Cárdenas et al. For 60 bp read lengths, *Centrifuge* has a higher sensitivity (91.3%) compared to *Kraken2* (88.2%) and precision (94.0%) compared to *Kraken2* (92.3%). Generally, both models demonstrate improved performance with longer reads, though their sensitivity and precision tend to plateau beyond 80 bp. Both *Centrifuge* and *Kraken2* demonstrate robust performance in classifying deaminated reads, with only a slight decrease in mean sensitivity from 91.7% to 90.0% for *Centrifuge* and 87.6% to 87.0% for Kraken2 when single-strand deamination probability was increased from 0% to 50% (**Figure S1C**). We observe almost no change in precision across the range of deamination probabilities (*Centrifuge* 94.05% to 93.49%, *Kraken2* 92.27% to 92.15%) (**Figure S1D**).

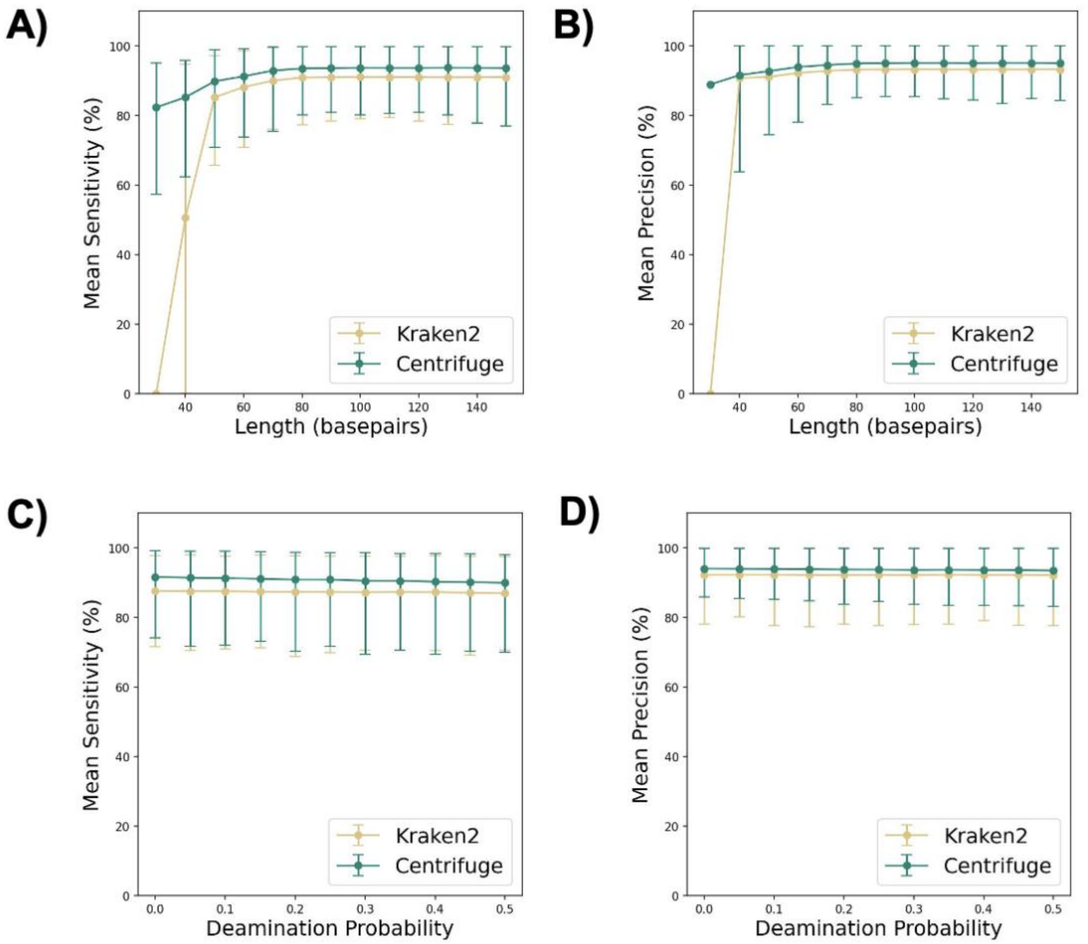


**Figure S1. Precision and sensitivity for the correct identification of viral reads by *Centrifuge* and *Kraken2*.** Sensitivity is defined as the number of reads correctly recovered at a species level divided by the total number of reads simulated. Precision is defined as the number of reads correctly recovered at a species level divided by the total number of reads recovered. Mean **A)** sensitivity and **B)** precision of 238 viral genomes, with read length from 30 bp to 150 bp with 10 bp increments. Error bar shows 10th percentile to 90th percentile. Mean **C)** sensitivity and **D)** precision in relation to single-strand deamination probability increasing from 0% to 50% with 5% increments. Error bar shows 10th percentile to 90th percentile.

*Metagenomic ancient hominin simulations*

To determine if the above results are applicable to the detection of ancient hominin sequences, we created three different simulation datasets (**Supplementary Table 3**), each with increasing complexity. In total 10 million reads were simulated for each dataset with lengths from 20 to 100 bp. Double-stranded deamination was added during Neanderthal read simulation assuming the Briggs model[27] with an average length of 0.4 of single-stranded overhanging ends, 3% single-stranded nick frequency, 30% deaminated cytosine residuals in single-stranded DNA, and 1% deaminated in double-stranded DNA.

**Supplementary Table 3:** Genetic data used to create simulated sequencing reads with *Gargammel*. For hominin data, alignments to the hg19 human reference genome were used.

| Simulation Dataset | Genome and composition | Relevant Reference(s) |
|---|---|---|
| Set 1 | 90% *Homo sapien* (NA12778 from 1000 genomes dataset, Chromosome 1) | [17] |
| | 10% Neanderthal (Altai, chromosome 22) | [28] |
| Set 2 | 40% *Homo sapien* (NA12778 from 1000 genomes dataset, Chromosome 1) | [17] |
| | 10% Neanderthal (Altai, chromosome 22) | [28] |
| | 50% Bacteria | [22,29] |
| Set 3 | 10% *Homo sapien* (NA12778 from 1000 genomes dataset, Chromosome 1) | [17] |
| | 1% Neanderthal (Altai, chromosome 22) | [28] |
| | 44.5% Bacteria | [22,29] |
| | 4.45% Bos Taurus (GCF_000003055.6) | [30] |
| | 4.45% Sus scrofa (GCA_001292865.1) | |
| | 6.675% Canis Lupus (GCA_905319855.2) | |
| | 6.675% Elephas maximus (GCA_014332765.1) | |
| | 13.35% Hyaena hyaena (GCA_004023945.1) | |
| | 8.9% Ursus americanus (GCA_003344425.1) | |

For these tests we used the pre-built full nucleotide reference databases for *Kraken2* (710GB, version from November 29, 2023) and *Centrifuge* (64GB, version from March 2018), each of which were downloaded from index zone on AWS (**Supplementary Table 1**). Sensitivity is defined as the number of reads correctly recovered at a species level divided by the total number of reads simulated that passed the minimum length cutoff, while the definition of precision remains the same as previously described.

We found that *Centrifuge* showed much higher sensitivity across all three simulation datasets, capturing at least 18% more reads. In the most complex scenario, *Centrifuge* did show reduced precision compared to *Kraken2* (97.5% compared to 99.75% for Set 3) (**Figure S2A**). The memory requirement difference however was drastic, where our test used 139GB of maximum RSS memory with *Centrifuge*, compared to 710GB with *Kraken2*, mainly due to the size of the pre-built index database (**Figure S2B**). The overall run time was on the same order of magnitude, but *Centrifuge* ran 4 times faster in Set 3 when the samples consisted of multiple species (**Figure S2C**).

Given the lower memory requirement and higher sensitivity, we recommend the use of *Centrifuge*, which is our default classification model for *sedimix*. All further analysis below, if not specifically mentioned, uses *Centrifuge* for the classification step.

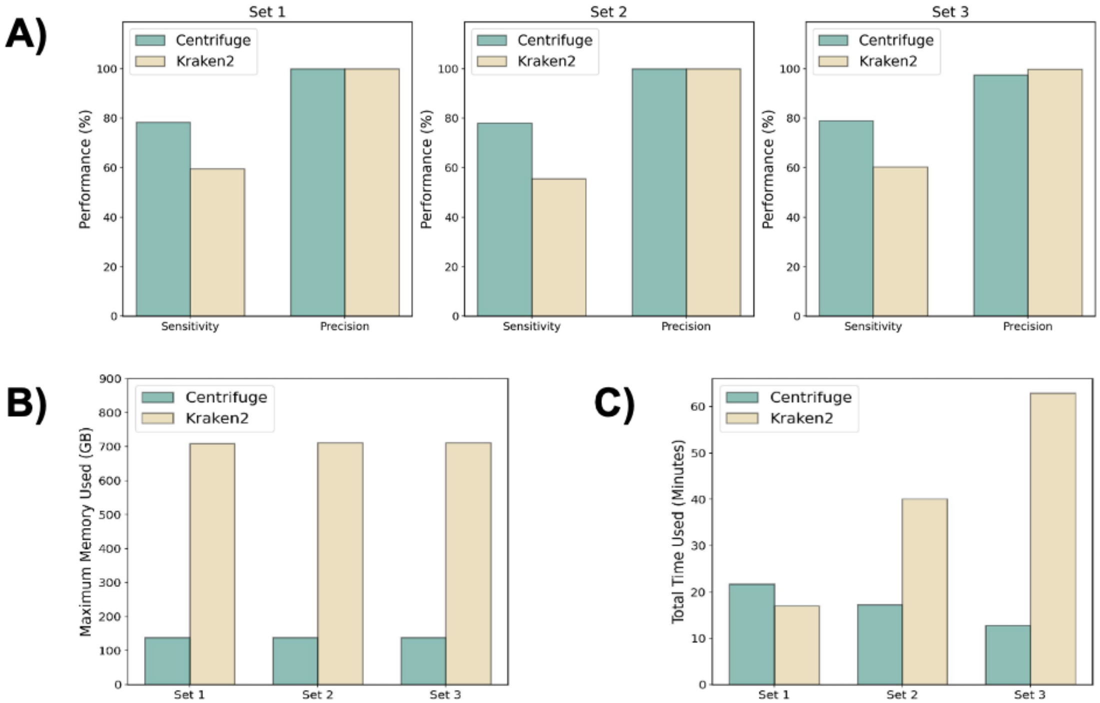


**Figure S2. Performance of *Centrifuge* and *Kraken2* based on three simulated hominin datasets. A)** Sensitivity and precision of each classifier across three simulated datasets. **B)** Memory usage and **C)** run time of each classifier across the different datasets. Memory usage is measured in maximum resident set size (RSS), which is roughly the largest total amount of physical memory assigned to a process in the workflow.

#### 3.2 Classification and mapping order

While it has been shown that combining classification and mapping can reduce exogenous contamination[31], the order of these steps has been used interchangeably. In Vernot et al.[3], mapping is performed first, followed by classification using *Kraken*. In contrast, Gelebret et al.[1] performs classification with *Centrifuge* followed by competitive mapping.

Technically, if the end goal is to identify hominin reads, the order of the pipeline should not matter, but it may impact the speed. Classification software has been shown overall to run at least one order of magnitude faster than mapping[32]. As sequencing data from sediments often includes tens of millions if not billions of reads, an order of magnitude difference would largely cut the run time from a few days to several hours. Therefore, we tested the impact of the ordering of these two steps with respect to sensitivity, precision, memory used, and run time.

We found that performing classification first largely reduces the number of reads and thus cutting the most time-consuming *bwa* mapping step. Performing classification first is almost 9 times faster compared to mapping first, processing the same 100 million reads in 45 minutes compared to 380 minutes (**Figure S3A**). As loading a large pre-built index database is the main limiting part for memory, memory requirement has less than a 17% difference (<20GB) between classification first and mapping first (**Figure S3B**). The minor differences observed in relation to sensitivity (**Figure S3C**) is due to the duplicate removal step in *sedimix*, which occurs directly after the mapping step.

An additional benefit of performing classification before mapping is that users are then provided with a general landscape of a sample's species composition. This can be helpful for further exploration of the non-hominin sequences. Thus, we recommend performing classification before mapping in the *sedimix* workflow due to the much faster run time and more informative intermediate outputs.

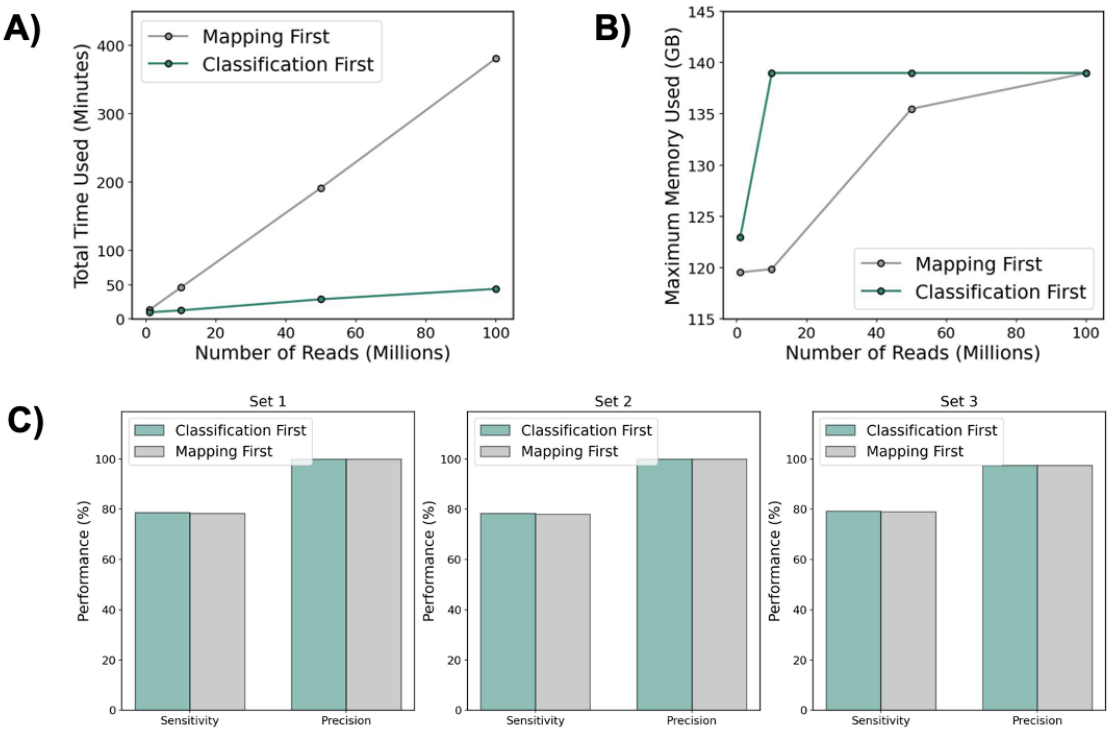


**Figure S3. Performance of *sedimix* in mapping-first or classification-first approach based on three simulated hominin datasets. A)** Run time and **B)** memory usage of either approach with increasing numbers of input reads. Datasets simulated under 95% bacteria and 5% Neanderthals for 1, 10, 50, and 100 million reads. Memory usage is measured in maximum RSS same as before. **C)** Sensitivity and precision of performing mapping-first or classification-first across three simulated datasets.

#### 3.3 Classification filtering level

Both *Kraken2* and *Centrifuge* output taxonomic assignments to different levels in the tree (family, genre, species, etc) based on each input sequence's similarity to all reference genomes. The taxonomic level of identification will then impact the number of reads retained for downstream analyses. As previous workflows have selected classification at the order level (primate)[3] and species level (*Homo sapiens*)[1], we tested the impact of these two selections on precision and sensitivity of the classification.

To test this we ran *sedimix* (classification by *Centrifuge* first performed before mapping) on the three simulation datasets (**Supplementary Table 3**). We discovered that filtering reads to only *homo sapiens* or the primate lineage has minimal impact on the sensitivity or precision (**Figure S4**). As expected, using a higher taxonomic rank increases the sensitivity by about 2% across all three datasets. The precision is the same for the first two datasets, while for the most complex dataset the precision is slightly higher when filtering on the species level (97.97% compared to 97.50%).

We recommend filtering at the “primate”’ level for the higher sensitivity and negligible difference in precision, though overall the difference in output is likely to be very small. If users wish to change the taxonomic level they would like to filter on, this can be specified in the input parameter file.

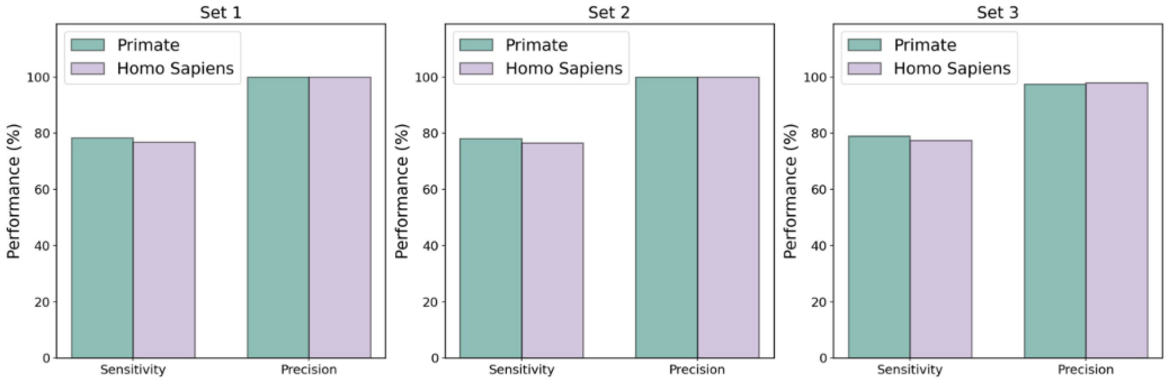


**Figure S4. *sedimix* performance when classification outputs are filtered on different taxonomic levels.**

### 3.4 Database complexity

In our previous testing, we found that loading the reference database by each classifier leads to high memory requirements (**Figure S3A**). There is a “standard” reference database build that is smaller, containing genomes from bacteria, archaea, viruses, and a human. We compared sensitivity, precision, run time, and memory requirements when running *sedimix* with this smaller database compared to the complete NCBI nucleotide database. We found that the smaller database had high sensitivity (82.4% compared to 79.0% for Set 3), but lower precision (96.1% compared to 97.5% for Set 3) for the identification of hominin reads (**Figure S5A**). This is expected as the increased granularity of the classification index will lead to non-hominin mammalian reads being correctly classified once these genomes are included in the reference database. The maximum memory requirement, as expected, is much lower when using the smaller database, though the usage varies with Set 3 using 35 GB of RAM while Set 2 uses 13 GB (**Figure S5B**). Run time was 2-4 times faster when using a smaller database, largely due to the time to load the index onto the server.

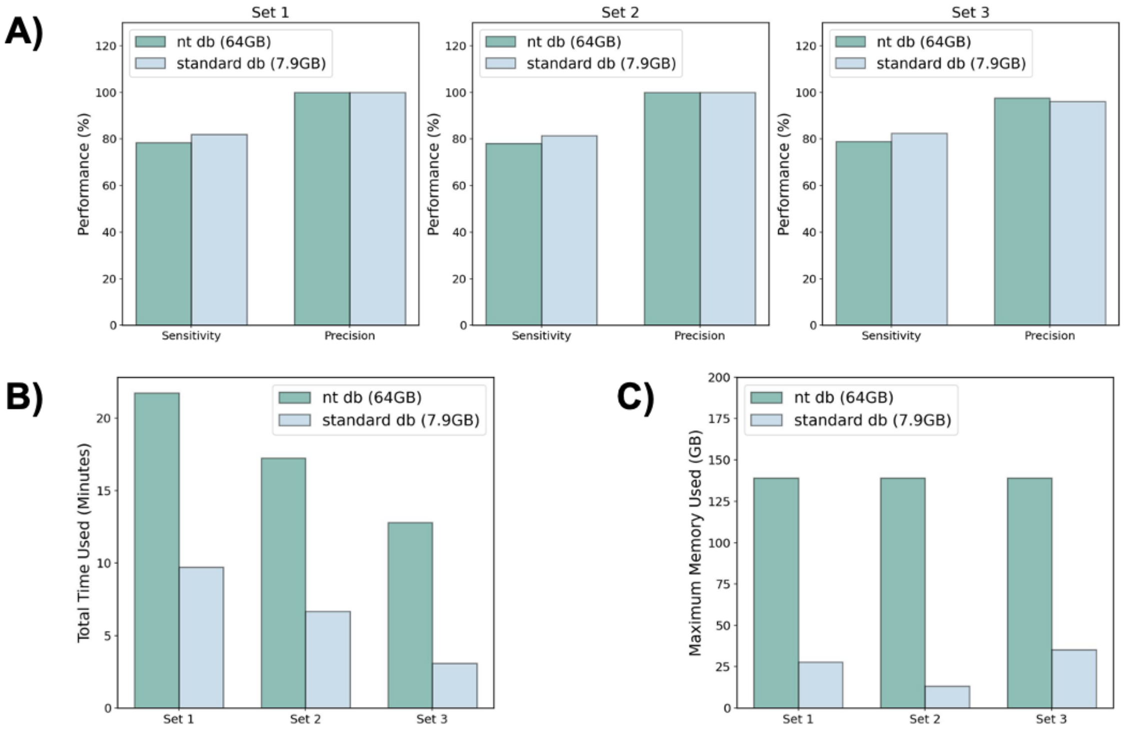


**Figure S5. Performance of *sedimix* in using two different sized classification reference databases on three simulated hominin datasets. A)** Sensitivity and precision of performing mapping-first or classification-first across three simulated datasets. **B)** Run time and **C)** memory usage of each reference database. Memory usage is measured in maximum RSS, the same as before.

We note that while there is little difference in the identification of hominin reads, only the larger reference database includes mammals. By looking at the intermediate files after classification in the most complex simulation (Set 3), we found that when using the smaller reference database we were unable to correctly identify the sequences deriving from mammals (**Figure S6**). Given that our simulation is only a simplification of real-world scenarios, we recommend larger, more complete databases to be used as default for *sedimix* to achieve a higher precision and improved classification of non-hominin compositions.

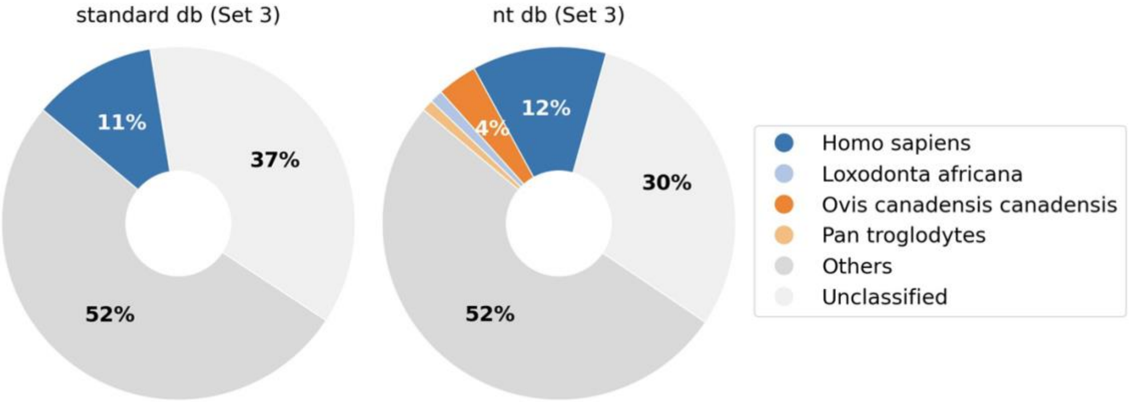

**Figure S6. Classifications of simulated reads with *Centrifuge* in using two different sized classification reference databases on Set 3.** Microbiome includes all species in bacteria, viruses, phages, viroids. Others include all species excluding microbiomes and the ones shown with labels.

### 4. Testing on Published Real Data

We tested *sedimix*'s performance on data from two previously published studies: (1) Gelabret et al. 2021 that includes ancient *hominin* DNA identified in a sediment sample[1] and (2) Essel et al. *2023* that includes touch DNA from an ancient *hominin* recovered from a cervid tooth pendant[2]. For each study, the data was downloaded from the European Nucleotide Archive (ENA) (project ID's PRJEB41420 and PRJEB56213 respectively) and then analyzed using our *sedimix* workflow.

#### 4.1 Comparison to Gelabert et al., 2021

*Gelabret et al* utilized shotgun sequencing to retrieve human and faunal environmental genomes from a 25,000-year-old Upper Paleolithic sediment sample from the Southern Caucasus[1]. We focused on the sample SAT29, which was identified as containing the highest number of reads mapping to the human nuclear genome and was used for downstream population genetics analyses. Briefly, after adapter trimming, in Gelabret et al. the authors removed low quality reads that contain at least 75% of bases with mapping quality scores below 30. Reads shorter than 30bp were also discarded after trimming two bp from both ends. The authors applied*Centrifuge* followed by competitive mapping to identify sequences related to *Homo sapiens* and other species. To replicate the results as closely as possible, we followed a similar filtering criteria, restricting the analysis to reads with minimum read length of 30 and a minimum mapping quality of 25, and then ran *sedimix* (without a SNP panel) with default parameters. We note the duplicate removal criterion is different between *sedimix* and the original paper, as the authors used *picard 2.21.4[33]* while we use *samtools markdup[9,10]*.

There were in total 522,997,582 reads in the raw sequencing file downloaded from ENA, which was slightly lower (6.8%) than the 561,263,536 reads reported in *Gelabret et al*. To compare the outputs of the two pipelines, we examined (1) percentage and count of reads assigned to hominin lineage, (2) reads identified as hominin sequences post de-duplication, (3) frequency of deamination on the 5' and 3' end of sequences. (**Supplementary Table 4**). *sedimix* assigned 0.096% (500,929) of the reads to hominin lineage, which is similar to 0.118% (661,765) inferred by Gellabert et al. 2021. To examine if this difference stems from differences in identifying duplicate reads (*picard* vs. *samtools*), we ran the same approach (*samtools markdup*) on the bam file generated by both pipelines. After duplicate filtering, we identify 0.094% (525,505) unique hominin sequences in Gellabert et al. 2021 data, while *sedimix* results in 0.096% (500,929) sequences. Mean fragment length was almost identical (46.2 bp compared to 46.3 bp).

We also compared the overlap between these sequences mapped by *sedimix* (500,929) and and found that ~74% of sequences (437,050) exist in both outputs, and 88,455 sequences (15%) exist only in Gelabert et al., while 63,879 (11%) sequences exist only in our pipeline (**Figure S7A**). Of those 88,455 sequences that we missed, we found that only 12,108 of these were classified as 'primate' by *Centrifuge* and the remaining 69,695 were labeled as 'unclassified'. For the 12,108 sequences assigned to primate, all are mapped, but they did not pass the *sedimix* mapping quality filter of 25 after *bwa aln*. While the classification method and nucleotide database used in our study and Gellabert et al. is similar, we cannot rule out some differences due to versions of the programs, databases as well as the human reference genome sequence used in the analysis.

**Supplementary Table 4: Comparison between *sedimix*'s output and Gelabert et al. original results.**

| | **Sample ID** | **Input file sequences** | **Percentage Identified hominin sequences [%] (count)** | **Percentage Identified hominin sequences after *samtools markdup* [%] *(count)*** | **Mean fragment length (bp)** | **5' C-to-T substitution frequency [%] (CI)** | **3' C-to-T substitution frequency [%] (CI)** |
|---|---|---|---|---|---|---|---|
| Original paper | SAT29 | 561,263,536 | 0.118 (661,765) | 0.094 (525,505) | 46.2 | 21 | 20 |
| *sedimix* | SAT29 | 522,997,582 | 0.096 (500,929) | 0.096 (500,929) | 46.3 | 20.1 (19.8-20.3) | 19.2 (19.0-19.5) |

In addition to the number of sequences recovered, we also examined the observed deamination of the recovered *homo sapiens* sequences. *sedimix*'s output has 5' deamination of 20.1% (95% CI: 19.8%-20.3%) and 3' deamination of 19.2% (95% CI: 19.0%-19.5%) (**Figure S7B**), which is very close to the original paper result with 5' deamination of 21% and 3' deamination of 20%.

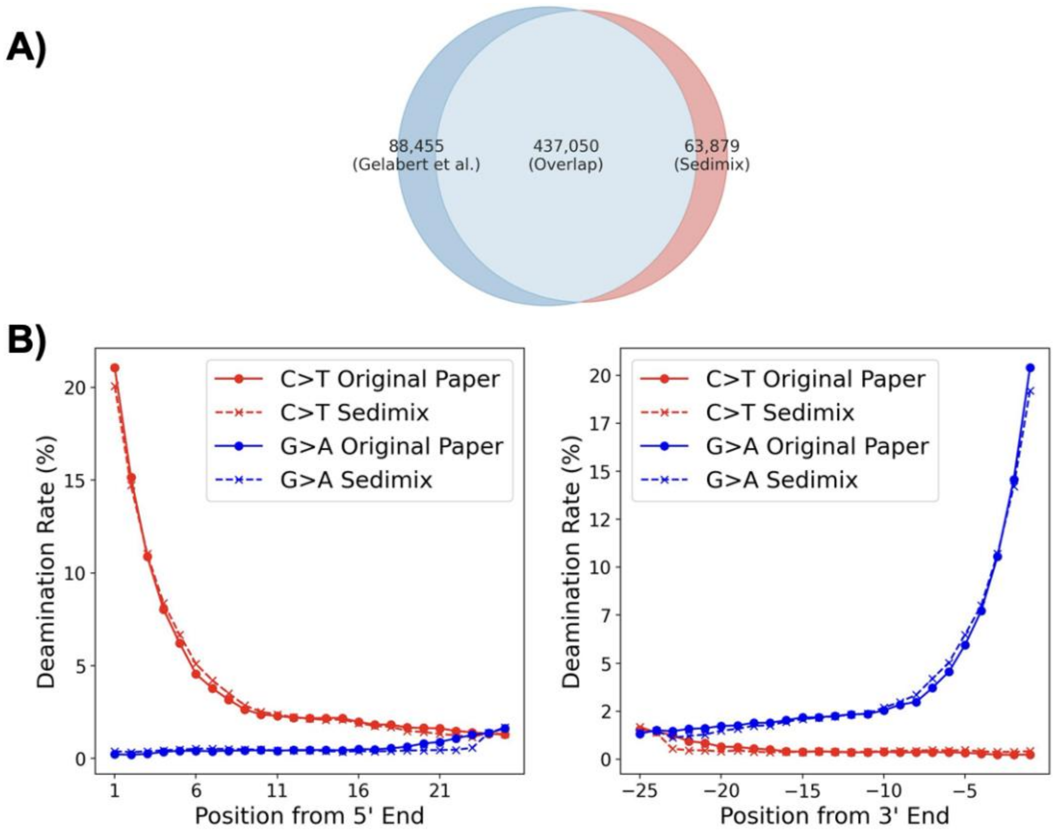


**Figure S7. Comparison between *sedimix* output and Gelabert et al. original results. A)** Overlap between the sequences identified in the paper and in *sedimix*.. **B)** Deamination patterns for the sequences identified in the paper and in *sedimix*. Both deamination profiles were generated by *mapDamage2*.

#### 4.2 Comparison to Essel et al., 2023

Essel et al. 2023 sequenced touch human nuclear DNA from an Upper Palaeolithic deer tooth pendant that was found in Denisova Cave[2]. This study utilized a designed capture probe named AA204[3] containing 470,724 SNPs. We downloaded this SNP file and built the alternative reference genome by replacing the allele at each SNP into a third allele that is neither the alternative nor the reference allele at that position, and then ran *sedimix* parameter *use_snp_panel* to True.

In our comparison, we focused on three published libraries, including N12015, N12018, and N12021. The publically available data from ENA contained merged data from multiple libraries. Therefore we limited our comparisons to the published summary statistics instead of comparing the raw sequence counts directly. We applied *sedimix* to these data and identified hominin DNA reads. We recovered around 20% more reads across all three libraries compared to Essel et al. 2023. The number of SNPs recovered using *sedimix* is also 30% higher for all three libraries. The proportion of hominin fragments we estimate is very similar to Essel et al. with differences within

1%. Similarly, the 5’ C-to-T deamination rate and 3’ C-to-T deamination rate are also qualitatively similar (with overlapping confidence intervals)(**Supplementary Table 5**).

**Table S5. Comparison between *sedimix*’s output and Essel et al. original results**. We note the discrepancy in the count of sequences generated stem from a minor error in the original paper and the numbers reported here for *sedimix* are correct (Elena Zavala, personal communication on behalf of the authors of the original paper).

| | **Captured Library ID** | **Sequences generated** | **Percentage of hominin fragments [%]** | **Unique on-target sequences** | **Number of SNPs covered** | **5' C-to-T substitution frequency [%] (95% conf. int.)** | **3' C-to-T substitution frequency [%] (95% conf. int.)** |
|---|---|---|---|---|---|---|---|
| Original paper | N12015 | 8,500,608 | 99.9 | 102,313 | 80,536 | 27.4 (26.9-28.0) | 18.0 (17.5-18.5) |
| | N12018 | 10,912,518 | 99.9 | 140,668 | 106,473 | 27.3 (26.8-27.7) | 18.0 (17.5-18.4) |
| | N12021 | 11,226,057 | 99.9 | 121,200 | 94,206 | 27.4 (26.9-27.9) | 18.1 (17.7-18.6) |
| *sedimix* | N12015 | 19,682,462 | 99.1 | 122,640 | 105,503 | 26.2 (25.7-26.7) | 17.0 (16.6-17.5) |
| | N12018 | 23,718,649 | 98.8 | 167,238 | 137,319 | 26.1 (25.7-26.5) | 16.9 (16.6-17.3) |
| | N12021 | 24,585,017 | 98.9 | 144,315 | 121,697 | 26.2 (25.7-26.6) | 17.1 (16.7-17.5) |

Overall, we have shown that *sedimix* produces reliable results for both real data examples, obtaining consistent results with published estimates. We provide an open-source snakemake pipeline, *sedimix,* to aid researchers in the analysis of ancient hominin DNA from sediments and increase reproducibility and ease of comparisons between studies.

### References:


1. Gelabert P, Sawyer S, Bergström A *et al.* Genome-scale sequencing and analysis of human, wolf, and bison DNA from 25,000-year-old sediment. *Curr Biol* 2021;**31**:3564–74.e9.

2. Essel E, Zavala EI, Schulz-Kornas E *et al.* Ancient human DNA recovered from a Palaeolithic pendant. *Nature* 2023;**618**:328–32.

3. Vernot B, Zavala EI, Gómez-Olivencia A *et al.* Unearthing Neanderthal population history using nuclear and mitochondrial DNA from cave sediments. *Science* 2021;**372**, DOI: 10.1126/science.abf1667.

4. Günther T, Nettelblad C. The presence and impact of reference bias on population genomic studies of prehistoric human populations. *PLoS Genet* 2019;**15**:e1008302.

5. Kim D, Song L, Breitwieser FP *et al.* Centrifuge: rapid and sensitive classification of metagenomic sequences. *Genome Res* 2016;**26**:1721–9.

6. Wood DE, Lu J, Langmead B. Improved metagenomic analysis with Kraken 2. *Genome Biol* 2019;**20**:257.

7. Li H. seqtk Toolkit for processing sequences in FASTA/Q formats. *GitHub*.

8. Li H, Durbin R. Fast and accurate short read alignment with Burrows–Wheeler transform. *Bioinformatics* 2009;**25**:1754–60.

9. Danecek P, Bonfield JK, Liddle J *et al.* Twelve years of SAMtools and BCFtools. *Gigascience* 2021;**10**, DOI: 10.1093/gigascience/giab008.

10. Bonfield JK, Marshall J, Danecek P *et al.* HTSlib: C library for reading/writing high-throughput sequencing data. *Gigascience* 2021;**10**, DOI: 10.1093/gigascience/giab007.

11. Quinlan AR, Hall IM. BEDTools: a flexible suite of utilities for comparing genomic features. *Bioinformatics* 2010;**26**:841–2.

12. Jónsson H, Ginolhac A, Schubert M *et al.* mapDamage2.0: fast approximate Bayesian estimates of ancient DNA damage parameters. *Bioinformatics* 2013;**29**:1682–4.

13. Perez G, Barber GP, Benet-Pages A *et al.* The UCSC Genome Browser database: 2025 update. *Nucleic Acids Res* 2025;**53**:D1243–9.

14. Smith TF, Waterman MS, Burks C. The statistical distribution of nucleic acid similarities. *Nucleic Acids Res* 1985;**13**:645–56.

15. de Filippo C, Meyer M, Prüfer K. Quantifying and reducing spurious alignments for the analysis of ultra-short ancient DNA sequences. *BMC Biol* 2018;**16**:121.

16. Prüfer K. Alignments of modern and archaic human genomes with primate outgroup genomes in tab-separated-values format. 2021, DOI: 10.17617/3.5H.

17. 1000 Genomes Project Consortium, Auton A, Brooks LD *et al.* A global reference for human genetic variation. *Nature* 2015;**526**:68–74.

18. Wood DE, Salzberg SL. Kraken: ultrafast metagenomic sequence classification using exact alignments. *Genome Biol* 2014;**15**:R46.

19. Arizmendi Cárdenas YO, Neuenschwander S, Malaspinas A-S. Benchmarking metagenomics classifiers on ancient viral DNA: a simulation study. *PeerJ* 2022;**10**:e12784.

20. Buchfink B, Reuter K, Drost H-G. Sensitive protein alignments at tree-of-life scale using DIAMOND. *Nat Methods* 2021;**18**:366–8.

21. Truong DT, Franzosa EA, Tickle TL *et al.* MetaPhlAn2 for enhanced metagenomic taxonomic profiling. *Nat Methods* 2015;**12**:902–3.

22. Renaud G, Hanghøj K, Willerslev E *et al.* gargammel: a sequence simulator for ancient DNA. *Bioinformatics* 2017;**33**:577–9.

23. Mihara T, Nishimura Y, Shimizu Y *et al.* Linking virus genomes with host taxonomy. *Viruses* 2016;**8**:66.

24. Brister JR, Ako-Adjei D, Bao Y *et al.* NCBI viral genomes resource. *Nucleic Acids Res* 2015;**43**:D571–7.

25. O'Leary NA, Wright MW, Brister JR *et al.* Reference sequence (RefSeq) database at NCBI: current status, taxonomic expansion, and functional annotation. *Nucleic Acids Res* 2016;**44**:D733–45.

26. Turner G, Barbulescu M, Su M *et al.* Insertional polymorphisms of full-length endogenous retroviruses in humans. *Curr Biol* 2001;**11**:1531–5.

27. Briggs AW, Stenzel U, Johnson PLF *et al.* Patterns of damage in genomic DNA sequences from a Neandertal. *Proc Natl Acad Sci U S A* 2007;**104**:14616–21.

28. Prüfer K, Racimo F, Patterson N *et al.* The complete genome sequence of a Neanderthal from the Altai Mountains. *Nature* 2014;**505**:43–9.

29. Seguin-Orlando A, Korneliussen TS, Sikora M *et al.* Paleogenomics. Genomic structure in Europeans dating back at least 36,200 years. *Science* 2014;**346**:1113–8.

30. The Bovine Genome Sequencing and Analysis Consortium, Elsik CG, Tellam RL *et al.* The genome sequence of taurine cattle: A window to ruminant biology and evolution. *Science* 2009;**324**:522–8.

31. Ravishankar S, Perez V, Davidson R *et al.* Filtering out the noise: metagenomic classifiers optimize ancient DNA mapping. *Brief Bioinform* 2024;**26**, DOI: 10.1093/bib/bbae646.

32. Jurado-Rueda F, Alonso-Guirado L, Perea-Cham-Blee TE *et al.* Benchmarking of microbiome detection tools on RNA-seq synthetic databases according to diverse conditions. *Bioinform Adv* 2023;**3**:vbad014.

33. Broad Institute. Picard Toolkit. *Broad Institute, GitHub Repository* 2018.
